# MINFLUX Reveals Dynein Stepping in Live Neurons

**DOI:** 10.1101/2024.05.22.595351

**Authors:** Jonas M. Schleske, Jasmine Hubrich, Jan Otto Wirth, Elisa D’Este, Johann Engelhardt, Stefan W. Hell

## Abstract

Dynein is the primary molecular motor responsible for retrograde intracellular transport of a variety of cargoes, performing successive nanometer-sized steps within milliseconds. Due to the limited spatiotemporal precision of established methods for molecular tracking, current knowledge of dynein stepping is essentially limited to slowed-down measurements in vitro. Here, we use MINFLUX fluorophore localization to directly track CRISPR/Cas9-tagged endogenous dynein with nanometer/millisecond precision in living primary neurons. We show that endogenous dynein primarily takes 8 nm steps, including frequent sideways steps but few backward steps. Strikingly, the majority of direction reversals between retrograde and anterograde movement occurred on the time scale of single steps (16 ms), suggesting a rapid regulatory reversal mechanism. Tug-of-war-like behavior during pauses or reversals was unexpectedly rare. By analyzing the dwell time between steps, we concluded that a single rate-limiting process underlies the dynein stepping mechanism whereby dynein consumes one adenosine 5’-triphosphate (ATP) per step. Our study underscores the power of MINFLUX localization to elucidate the spatiotemporal changes underlying protein function in living cells.

---

Axons are the longest processes of neurons, reaching lengths of millimeters to meters. For rapid and targeted movement of cellular components such as mitochondria, organelles, proteins or RNA between the soma and the axon terminal, a directed, non-diffusive transport mechanism is essential (1, 2). Due to the crucial role of intracellular transport in neuronal development and cellular integrity, several neurological diseases are associated with axonal transport impairment (3, 4). Axonal transport is facilitated by the kinesin motor protein family and cytoplasmic dynein 1 (hereafter dynein), which move in opposite directions on the unidirectionally oriented microtubules in the axon (5-7). For retrograde transport, one or two dynein dimers are assembled with dynactin and a specific activating adaptor tethered to the cargo (8-11).

Dynein has been studied extensively over the past two decades. The use of artificially dimerized, truncated yeast dynein monomers in vitro revealed that dynein moves towards the minus end of microtubules in steps of high variability (12, 13), relying on a proposed flexible and uncoordinated stepping mechanism (13-15). Furthermore, highly variable step sizes between 8 and 32 nm, including frequent sideways and backward steps, were observed using purified human and mammalian dynein motor complexes (16, 17). Additionally, it was found that the dynein motor complexes take smaller steps, mainly 8 nm in size, under load. However, due to the limited spatiotemporal precision, either individual steps could not be resolved in live cells (18, 19), or drastically reduced adenosine 5′-triphosphate (ATP) concentrations were used to slow down the dynein motor (complex) movement and allow single step observation in vitro (12-17). As a result, the current understanding of the dynein stepping behavior is largely based on decidedly slowed-down in vitro movements, raising the question of how endogenous dynein actually steps in a living cell. Although cargo and large artificial cargo serving as optical probes have been tracked with high precision in living cells (20-22), it is challenging to extrapolate these observations to the dynein stepping itself. Direct tracking of endogenous dynein in living cells requires a method to resolve the dynein steps with a specific minimally invasive label that is substantially smaller than the protein itself.

Recently, MINFLUX has been demonstrated to be capable of directly tracking protein dynamics and conformational changes, such as those of kinesin-1 (23, 24), by the use of a minimally invasive fluorescent label. MINFLUX localizes a fluorophore by relating its unknown position to the known position of the central intensity minimum (zero) of a fluorescence excitation beam (25). Reducing the distance between the two positions, i.e. matching them as closely as possible, increases the localization precision per detected fluorescence photon. MINFLUX typically requires only about 100 detected photons to achieve single-digit nanometer localization precision, meaning that a spatiotemporal precision of a few nanometers per millisecond is readily achieved. In contrast, popular fluorophore localization methods that rely on the identification of the maximum of the fluorescence diffraction pattern produced by the fluorophore on a camera (i.e. centroid calculation) require about one hundred times more detected photons per fluorophore (26). Consequently, to achieve comparable spatiotemporal precision, established methods require photon-detection rates that are accordingly higher. In many cases, these rates can be provided only by strongly scattering labels, such as beads, that are many times larger than the protein to be observed.

Here, we use MINFLUX localization and a label that is considerably smaller than the protein under observation to directly track endogenous mammalian dynein with nanometer/millisecond precision in a primary culture of living hippocampal neurons. Using a CRISPR/Cas9-mediated knock-in approach, we were able to specifically and directly tag dynein at two different sites and label them with a fluorophore. Our observations revealed that endogenous tail-labeled dynein performs mainly 8 nm steps along the microtubule, as well as frequent sideways but few backward steps. Most notably, direction reversals between retrograde and anterograde movement occurred on the time scale of individual steps, suggesting a rapid regulatory reversal mechanism. The analysis of the dwell time between steps led us to conclude that a single rate-limiting process underlies the dynein stepping mechanism, indicating that dynein requires a single ATP molecule per step.

## Direct and site-specific CRISPR/Cas9-mediated endogenous tagging of mammalian dynein in live primary neurons

Overexpression of individual dynein subunits has been shown to affect the integrity of the entire motor complex and endomembrane localization (27, 28). To avoid overexpression, we used CRISPR/Cas9-mediated endogenous tagging to directly and specifically tag dynein for subsequent study of its stepping behavior (29). For each dynein-specific target sequence (Tab. S1), we designed and generated an ORANGE-based (30) plasmid for the knock-in of a HaloTag (31) at either the N-terminus of the dynein heavy chain (Halo-DHC) or the N-terminus of the dynein intermediate chain (Halo-DIC) (Fig. S1; Tab. S2). Correct and specific integration of the HaloTag sequence at the dynein loci was assessed by sequencing the isolated genomic DNA of primary rat hippocampal neurons electroporated with each plasmid.

To visualize endogenous dynein in living neurons, we electroporated primary rat hippocampal neurons with one of our CRISPR/Cas9 knock-in plasmids. After six to nine days in culture, we labeled Halo-DHC and Halo-DIC with MaP555-Halo (32), and performed widefield microscopy tracking of endogenous dynein on stained microtubules (Fig. 1A). Thus, the Halo-DHC and Halo-DIC positive neurons were readily distinguishable from the wildtype neurons, and the axon was clearly identifiable as the longest and smoothest process (Fig. 1B and D). We followed the axon for several hundred microns, recorded videos of labeled dynein, and generated kymographs of Halo-DHC (Fig. 1C) and Halo-DIC (Fig. 1E). To distinguish between retrograde and anterograde movements, we applied Fourier filtering to the kymographs (33). In accordance with a recent study of endogenously tagged dynein in induced neurons (34), the majority of dynein was not processive. Instead, only a small fraction showed retrograde movement, including direction reversals and pauses (Fig. 1C and E; Fig. S2; Video S1), as previously observed for various intracellular cargoes (21, 35-39). Both processive Halo-DHC and Halo-DIC tagged motor complexes moved similarly fast in the retrograde direction (DHC: (1.1 ± 0.4) µm s^-1^, DIC: (1.5 ± 0.7) µm s^-1^, median ± median absolute deviation, MAD) with speeds up to 4 µm s^-1^, comparable to speeds reported for endogenously tagged dynein in induced neurons (34), prion protein (37) or amyloid precursor protein vesicles (40). Limitations to widefield tracking of individual dynein motors were set by non-processive labeled dynein molecules, but also by photobleaching and out-of-focus movement. In any case, CRISPR/Cas9-mediated endogenous tagging turned out to be highly suitable for direct and site-specific labeling of dynein in living primary neurons.

**Figure 1.**
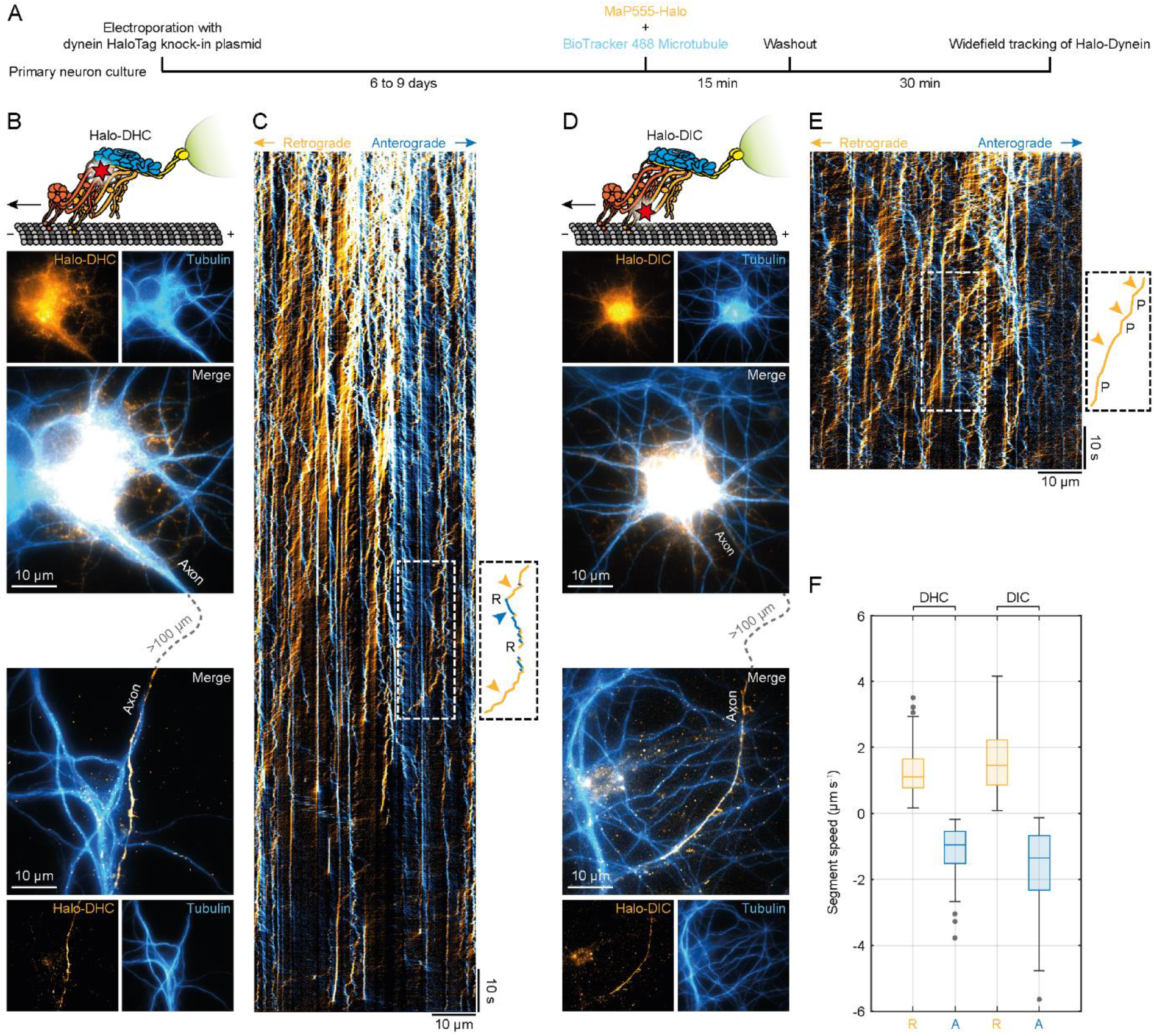
Direct and specific CRISPR/Cas9-mediated endogenous tagging of dynein in live primary neurons. (A) Schematic of sample preparation for widefield tracking of endogenous dynein (Halo-DHC or Halo-DIC). Primary rat hippocampal neurons were electroporated with a CRISPR/Cas9 knock-in plasmid, resulting in N-terminal tagging of endogenous dynein with a HaloTag. After six to nine days in culture, dynein and microtubules were labeled simultaneously with cell-permeable dyes (MaP555-Halo and BioTracker 488 Microtubule), washed, and imaged. (B) Sketch of the Halo-DHC motor complex labeled with MaP555-Halo (red star), including dynactin (blue), an activating adaptor (yellow), and a cargo (green). Example widefield images (maximum intensity projection of video) showing the soma and axon of a Halo-DHC positive neuron and its microtubules, as well as those of neighboring neurons. Videos of labeled dynein were recorded along the axon, which was identified as the longest and smoothest process and traced for several hundred microns. (C) Example kymograph of Halo-DHC along the axon with retrograde movement colored in orange and anterograde movement colored in blue. Overlapping motion appears in white. Inset shows initial retrograde movement, which is followed by an instantaneous direction reversal (R) to anterograde movement, and a prolonged reversal to retrograde movement, as indicated by colored arrowheads. (D) Sketch of the Halo-DIC motor complex and example widefield images showing the soma and axon of a Halo-DIC positive neuron. Videos were acquired similarly to Halo-DHC. (E) Example kymograph of Halo-DIC along the axon, where the inset shows a retrograde movement interrupted by pauses (P). (F) Boxplot of segment speed of retrograde and anterograde motion from Halo-DHC (n = 329, 25 videos, N = 6) and Halo-DIC (n = 349, 22 videos, N = 4).

## MINFLUX reveals stepping behavior of endogenous dynein

Next, we sought to investigate the stepping behavior of endogenous dynein. Again, we used our CRISPR/Cas9 knock-in plasmids to insert a HaloTag sequence into the dynein loci. To achieve single-molecule conditions and avoid having multiple labels per dynein motor complex, Halo-DHC or Halo-DIC were labeled with a low concentration of 100 pM JFX650-Halo (41). In a second step, the remaining dynein was saturated with MaP555-Halo (32) to identify Halo-Dynein positive neurons. Simultaneously, the microtubules of all neurons were labeled with BioTracker 488 to verify that MaP555-Halo was bound specifically to dynein on microtubules of only Halo-Dynein positive neurons (Fig. 2A).

**Figure 2.**
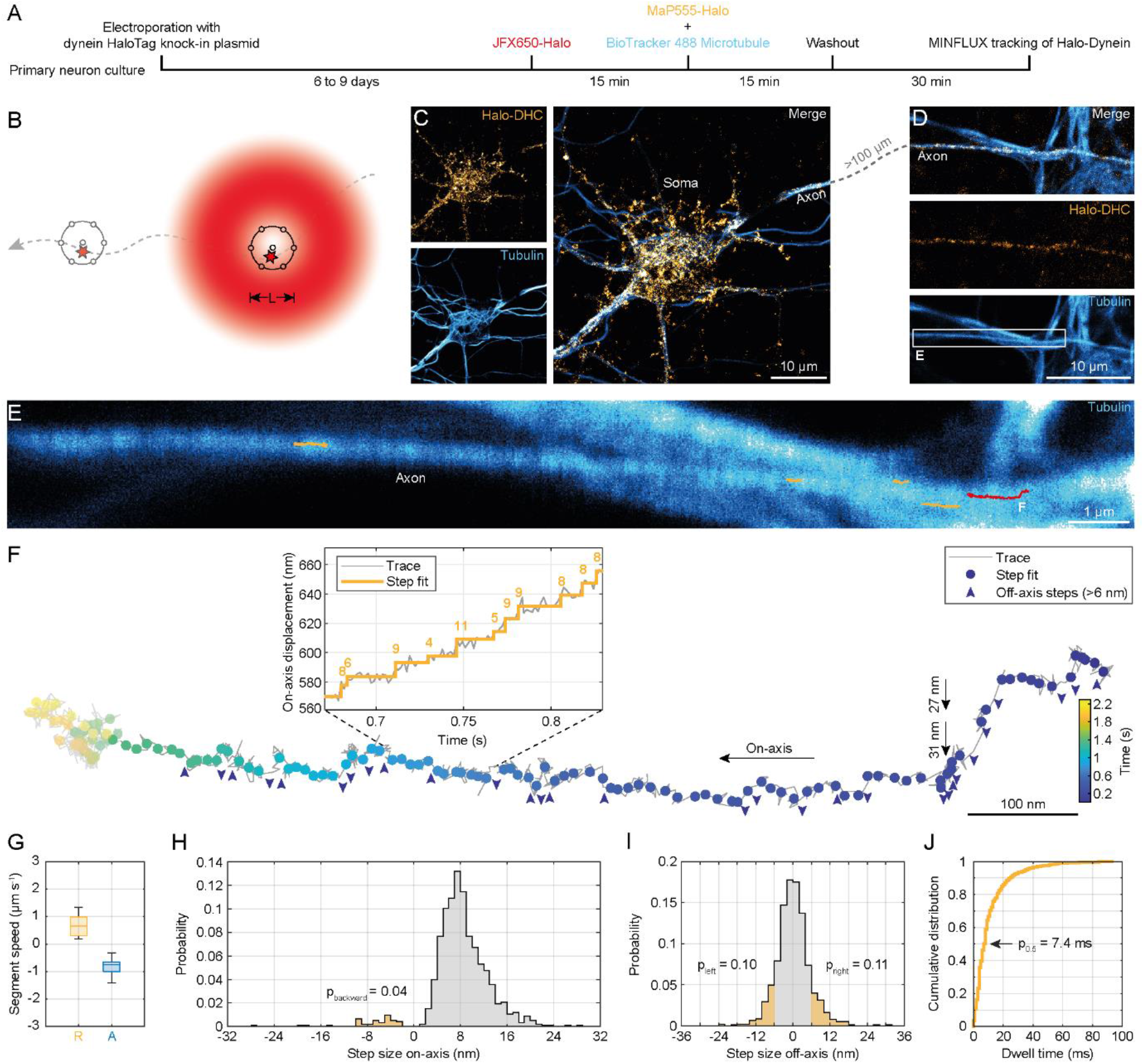
MINFLUX reveals stepping behavior of endogenous dynein in live neurons. (A) Schematic of sample preparation for MINFLUX tracking of endogenous dynein. Primary rat hippocampal neurons were electroporated with a CRISPR/Cas9 knock-in plasmid, resulting in N-terminal tagging of endogenous dynein with a HaloTag. After six to nine days in culture, a fraction of dynein was labeled with 100 pM JFX650-Halo. The remaining dynein was labeled with 100nM MaP555-Halo, and microtubules were simultaneously labeled. (B) MINFLUX procedure for tracking a single moving fluorophore (star on dashed trajectory). To determine the current position of the fluorophore with nanometer precision, a donut-shaped excitation beam (red) probes seven positions (black circles) around the fluorophore with diameter L. This procedure is repeated during the fluorophore movement. (C) Example confocal images showing a Halo-DHC positive neuron and microtubules. (D) Confocal images showing the axon of the Halo-DHC positive neuron whose axon was traced over several hundred microns. The selected MINFLUX region of interest is highlighted by the white box. (E) Overlay of confocal image of tubulin with five MINFLUX traces of Halo-DHC (orange and red) along the axon. (F) Example of a MINFLUX trace of Halo-DHC with two large off-axis steps (black arrows) indicating a change of multiple protofilaments or a sideways step to an adjacent microtubule. Dots colored by time show the steps of dynein. Off-axis steps larger than 6 nm are indicated by blue arrowheads. The step sizes are shown in the on-axis displacement versus time plot. The left end of this trace (grayed out) was not included in further analysis because there was no processive movement in this region. (G) Boxplot of segment speeds in the retrograde (n = 25) and anterograde directions (n = 17) of Halo-DHC. (H) On-axis step size distribution of Halo-DHC with indicated proportion of backward steps (n = 925, N = 6). (I) Off-axis step size distribution of Halo-DHC with highlighted proportions of off-axis steps larger than 6 nm in either direction (n = 925, N = 6). (J) Cumulative distribution of dwell times between consecutive steps of Halo-DHC (n = 900, N = 6).

To resolve individual steps of endogenous dynein, we used a MINFLUX microscope (42), whose nanometer/millisecond spatiotemporal precision is achieved by probing the current position of a single moving fluorophore with the intensity minimum of a donut-shaped excitation beam (Fig. 2B). To ensure that only one emitter is localized at a time, the ratio of central to peripheral emission was calculated during tracking as a termination condition of the localization scheme (42). Using the MaP555-labeled dynein, we identified a Halo-DHC positive neuron and followed its axon over several hundred microns (Fig. 2C). Note that tubulin staining reveals a neighboring wildtype neuron, yet without colocalizing MaP555 emission, confirming the specificity of the labeling (Fig. 2D). Using this procedure, we were able to record MINFLUX traces of Halo-DHC in living neurons up to 1.6 µm in length and 3 s in duration in the retrograde and anterograde directions (Fig. S3). We obtained a median localization precision of (3.5 ± 0.4) nm using a pattern diameter of L = 75 nm (Tab. S4), which is comparable to state-of-the-art MINFLUX tracking studies (23, 24). For the recorded MINFLUX traces of Halo-DHC along the axon (Fig. 2E and F), steps were identified using a bias-free step detection algorithm with no prior assumptions about step number or size (43), resulting in a median step size uncertainty of (2.1 ± 0.6) nm.

To quantitatively characterize the stepping behavior of endogenous dynein, we classified all MINFLUX traces into retrograde and anterograde segments. The speed of Halo-DHC in both directions was similar (Fig. 2G), and comparable to the widefield recordings (Fig. 1F). Examining the detected steps in retrograde segments, most on-axis steps are approximately 8 nm in size, but the N-terminus of the DHC (dynein tail) also takes larger steps up to 16 nm (Fig. 2H). Consistent with previous reports (12, 16), dynein frequently steps sideways (Fig. 2I), perhaps to circumvent obstacles on the track (44), or also as an inherent consequence of dynein’s long stalk domain and flexibility, which permits high variability in both on-axis and sideways steps. Occasionally, we also observed large sideways steps (>25 nm), suggesting a change to an adjacent microtubule or a step over several protofilaments at once (Fig. 2F). The distribution of dwell times between successive steps of Halo-DHC shows that the steps follow each other very quickly (half of the steps are faster than 7.4 ms) and that dynein dwells for less than 100 ms before stepping again (Fig. 2J).

A comparison of the stepping behavior of Halo-DIC to that of Halo-DHC reveals that the on-axis step size distribution of Halo-DIC (Fig. S4B) is broader than that of Halo-DHC (Fig. 2H), exhibiting slightly larger steps and more frequent backsteps. Similarly, the off-axis distribution of Halo-DIC (Fig. S4C) is also wider than that of Halo-DHC (Fig. 2I). These observations indicate that the DIC N-terminus exhibits greater flexibility than the dynein tail while moving. This observation is consistent with structural studies indicating that the DIC N-terminus is quite flexible (8, 45).

Overall, the on-axis step size distribution of the dynein tail (Fig. 2H) is substantially narrower with smaller steps (4 to 16 nm) than in purified dynein motor complexes without load (8 to 32 nm) (16, 17). However, the distribution shows similarities to the in vitro distribution of dynein under high load (16, 46), which seems reasonable since dynein is only active in live cells when bound to a cargo. The mean forward step size of (8.7 ± 3.8) nm (Tab. S5) is comparable to that observed for dynein under high load in vitro (∼10 nm) (16). This finding suggests that dynein exerts a high force under endogenous conditions, assuming a molecular gear mechanism that predicts a decreasing step size with increasing load (17). Surprisingly, there were notable discrepancies in the observed incidence of backward steps compared to previous in vitro studies (12, 14, 16). Our observations of the dynein tail revealed a lower incidence of backward steps (p_backward_ = 0.04, Fig. 2H) in contrast to in vitro studies without load (∼0.20) (12, 14, 16) and an even higher discrepancy when high load was applied (0.35 to 0.50) (16). This suggests that dynein is highly efficient in moving forward under endogenous conditions, and that purified dynein does not fully resemble the dynein motor complex in live neurons.

## Naturally occurring direction reversals usually follow a rapid mechanism distinct from a tug-of-war model

It is widely accepted that multiple motors of opposite polarity are both anchored to the same intracellular cargo (2, 47, 48). Therefore, not only unidirectional but also bidirectional cargo movement has often been observed (21, 37, 39, 40). However, it is unclear how cargo directionality is controlled, whether by a stochastic process, typically referred to as a tug-of-war between motors of opposite polarity (49-51), regulated by adaptor or microtubule-associated proteins (52-54), post-translational modifications of microtubules (5), or by separate transport of different parts of the motor complex (34). With MINFLUX, it is possible to observe not only a few steps of the cargo movement with high precision (21), but a single dynein itself over several hundred nanometers. This allowed us to distinguish between processive retrograde and anterograde movements and thus to observe direction reversals with single-step resolution.

To study direction reversals and pauses at a spatial and temporal scale that was previously hidden by the lack of spatiotemporal precision in live-cell observations (18, 34), we categorized all direction reversals and pauses between successive processive movements of both Halo-DHC and Halo-DIC traces (Fig. S5, see Methods for details). We observed a similarly high percentage of traces containing direction reversals (38.5% of traces), both anterograde to retrograde and retrograde to anterograde, as well as pauses (41.0% of traces) between processive anterograde or retrograde movements (Fig. 3A). As expected, this suggests that motors of opposite polarity both act on the same cargo, possibly to bypass obstacles or as a means of regulation to reach the desired cargo destination (47, 48, 55). However, we also observed a large fraction of highly processive traces with neither reversals nor pauses (46.2% of traces), which could indicate the transport of different types of cargo in uninterrupted traces compared to traces with reversals and pauses.

**Figure 3.**
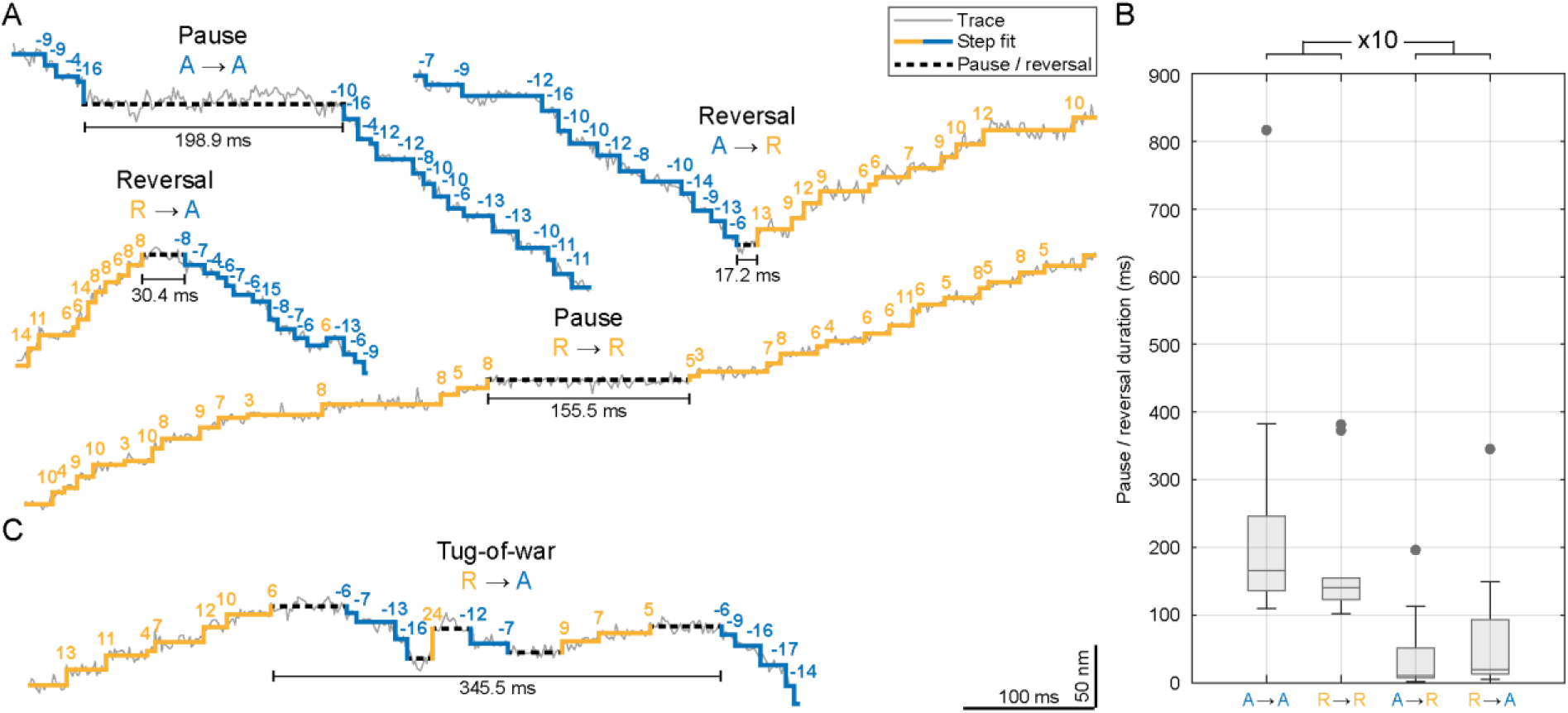
Direction reversals occur rapidly at the time scale of single steps. (A) Extracts from example MINFLUX traces of dynein containing either a pause between anterograde (A → A) or retrograde (R → R) processive movement or a directional reversal from anterograde to retrograde (A → R) or retrograde to anterograde (R → A). (B) Boxplot showing that reversals (n = 24) have durations on the order of dwell times of single steps and last ten times shorter than pauses (n = 25). Grey dots indicate outliers. (C) Extract from an example MINFLUX trace showing a potential tug-of-war between retrograde and anterograde movement.

Strikingly, direction reversals occurred on the order of the dwell times of single steps (Fig. 3B; see also Fig. 2J), suggesting a rapid regulatory reversal mechanism (37, 40, 56, 57). This rapid reversal is obviously in contrast to a tug-of-war scheme (50, 51, 58). Even more strikingly, pauses lasted substantially longer than single steps and were ∼10 times longer than the duration of direction reversals (Tab. S5). Unexpectedly, we rarely observed discrete forward and backward steps within reversals or pauses (Fig. 3C; 10.2% of reversals or pauses), which in turn seems consistent with a brief unregulated tug-of-war. Alternatively, this behavior could be the result of a slower regulatory mechanism that still allows for a brief mechanical competition of opposing motors (48). Altogether, we observed that rapid direction reversals are most common during axonal transport in living neurons, but in rare cases, motors (of opposite polarity) can still engage in a brief tug-of-war.

Compared to recent live-cell studies that also tracked dynein directly with conventional centroid localization (18, 34), the duration of reversals and pauses that we observed with MINFLUX was substantially shorter. This is because the spatiotemporal resolution of conventional tracking using single fluorophores or fluorescent proteins did not allow the detection of individual steps in living cells. Conversely, although we observed reversals and pauses of similar duration as previous widefield-based studies (Fig. 1C and E), and although we obtained MINFLUX traces over the course of several seconds, we were unable to characterize long-lasting reversals or pauses on the order of seconds, most likely due to photobleaching. Figure 2F shows an example of a MINFLUX trace in which dynein transitioned to a non-processive state at the end of the trace (grayed out area), but the attached fluorophore likely underwent photobleaching before we could track a potential resumption after a pause or reversal on the order of seconds. Nonetheless, the single-step precision of MINFLUX clearly allows characterization of naturally occurring direction reversals and pauses that were previously hidden, indicating a rapid regulatory reversal mechanism of bidirectional axonal transport in neurons.

## Dynein consumes one ATP per step

Whether dynein consumes one or more ATPs per step has remained controversial since the first structural studies, because dynein’s ring-shaped motor domain in principle allows ATP binding to four of its six ATPases associated with diverse cellular activities (AAA) domains (59). ATP hydrolysis at AAA1 has been shown to be essential for dynein motility (60). However, whether the AAA3 domain serves to regulate dynein (61, 62) or whether ATP hydrolysis at AAA3 must occur together with AAA1 to perform a single step, implying that dynein consumes two ATPs per step (20), remained the subject of debate.

To clarify whether dynein requires one or two ATPs per step, we performed a detailed analysis of the dwell times between consecutive steps of endogenous dynein. We compared three different models to describe the dwell time data, a single exponential decay and a convolution of two exponential decays with equal or unequal rate constants. To be independent of the data representation, we aimed to describe the data directly using maximum likelihood estimation. The model that best describes the data was determined using Akaike weights as a measure of the probability that one of the candidate models is the best, given the data and the three candidate models (63). According to this approach, the dwell time distribution was best described by a convolution of two exponentials with a low (*k*_1_) and a high (*k*_2_) rate constant (Fig. S6A and B). As our live-cell measurements were conducted at endogenous ATP concentrations and thus are not necessarily rate-limited by ATP binding, the lower rate constant can be assigned to the rate-limiting step of the ATPase cycle, while the higher rate constant may be attributed to fast cycle steps. Actually, the high rate constant is in good agreement with the ATP hydrolysis and ADP dissociation rate of ∼1000 s^-1^ determined in biochemical assays (64), whereas the low rate constant matches the ATPase activity of purified dynein-dynactin-adaptor complexes (16). However, as the extremely fast steps (τ < 3 ms) occur on a timescale close to the temporal resolution of our measurement, it is likely that they may be partially missed and thus underrepresented in the data. When considering only steps with dwell times τ < 3 ms, the single exponential model with a rate constant of *k* = (90 ± 3) s^−1^ provides the best fit to the data (Fig. 4A). Consequently, the low rate constant *k*_1_ of the model with different rate constants becomes equal to the rate constant *k* of the single exponential model.

**Figure 4.**
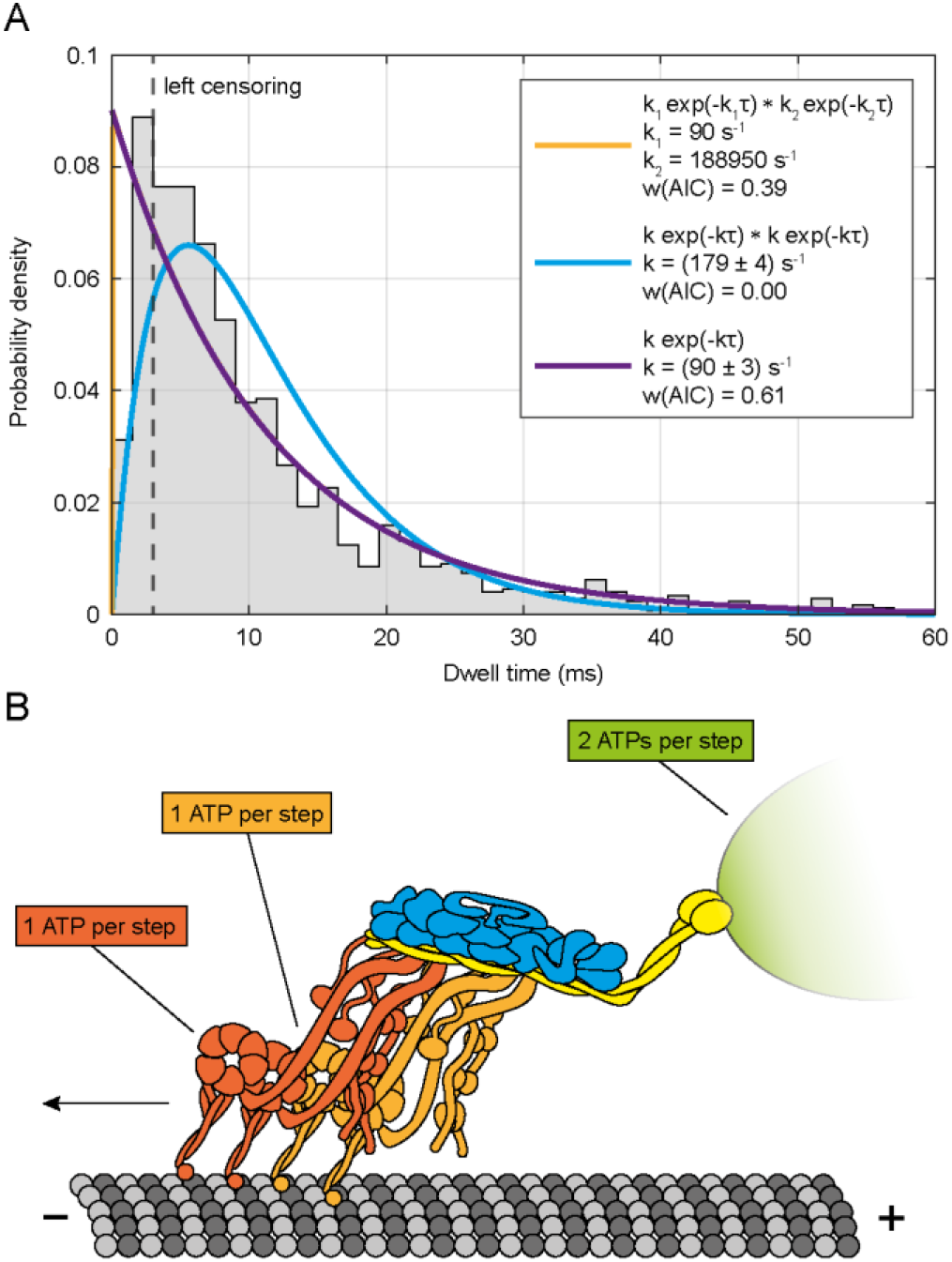
Dynein consumes one ATP per step. (A) Dwell time distribution of endogenous dynein (Halo-DHC and Halo-DIC) with three possible models for the description of the data (n = 1178, N = 23). Left censoring of the distribution corresponds to the case where it is assumed that extremely fast steps (τ < 3 ms) are partially missed and therefore underrepresented. Based on Akaike weights *w*(AIC), the single exponential model (*ke*^−*k*τ^, purple) is the favored model for the description of the data in this case (i.e. one ATP per step). The orange curve, representing a convolution of two exponentials with unequal rate constants 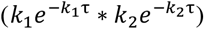, overlaps exactly with the purple curve, and the low rate constant *k*_1_ corresponds to that of the single exponential model, while the high rate constant *k*_2_ becomes infinite. A convolution of two exponentials with the same rate constant (*ke*^−*k*τ^ ∗ *ke*^−*k*τ^, cyan) would correspond to a model in which dynein consumes two ATPs per step, but does not describe the data well. 212 data points were excluded by the left censoring. (B) Model for the stepping behavior of endogenous dynein. Stepping of each dynein dimer (orange and red) is based on a single rate-limiting process and thus consumes one ATP per step. In the native case of two dynein dimers bound to dynactin (blue), constrained dynein heads might perform steps that only relieve intramolecular tension but do not contribute to the cargo (green) movement.

Both interpretations are consistent with different possible stepping mechanisms. One is an “alternating shuffle” (12), in which both head domains alternately take steps twice as large as the tail. The other is an “inchworm” model (15), in which one head domain takes the lead and the other head domain follows, which includes the possibility of uncoordinated head movement (13, 14). It is important to note that a convolution with two equal rate constants (i.e. two ATPs per step) does not adequately describe the data in either case. Instead, a single rate constant of ∼90 s^-1^ describes the ATPase cycle of dynein at the observed timescale. This implies that dynein consumes one ATP per step as proposed in in vitro studies (12, 16, 17, 46, 61), rather than two ATPs per step (20, 65).

When considered in conjunction with previous observations, our findings support a model for the stepping behavior of the dynein motor complex (Fig. 4B) that reconciles our observation in which dynein itself consumes only one ATP per step with the earlier observation that dynein-driven cargo movement requires two ATPs per step (20). Given that the native dynein motor complex is typically composed of two dynein dimers per dynactin (9, 34), the tight arrangement of the four dynein heads could imply that one head per dynein dimer is constrained in its movement (9, 45). When the unconstrained heads take steps, the constrained heads would have to catch up in order to relieve the intramolecular tension. However, the constrained heads are unable to contribute to cargo movement because they cannot pass the other heads. Such a scenario could result in a stepping mechanism consisting of two successive processes with the same rate constant (one step by the unconstrained head and one by the constrained head), with only the steps of the unconstrained heads contributing to cargo movement. Consequently, relying on two dynein dimers, cargo movement in a living cell would be best described by a convolution of two exponentials with equal rate constants (Fig. S6C), implying two ATPs per step for cargo movement, but one ATP per step for each dynein dimer.

In conclusion, MINFLUX localization enabled the direct tracking of CRISPR/Cas9-tagged endogenous dynein, revealing its stepping behavior in living primary neurons. We observed variable step sizes, down to 4 nm, of mammalian dynein moving on the microtubule lattice. Our high spatiotemporal precision enabled us to identify and characterize the fast changes in movement direction with an unprecedented level of detail, favoring a rapid regulated reversal mechanism over a stochastic tug-of-war. Finally, by analyzing the dwell times between steps, we conclude that dynein consumes one ATP per step, which clarifies previous findings. Our study illustrates the importance of combining direct and specific labeling of the endogenous protein of interest with the nanometer/millisecond tracking precision of MINFLUX for an accurate description of protein dynamics in living cells.

## Author contributions

J.M.S. designed and managed the dynein study, conducted the experiments, wrote the software to evaluate the MINFLUX traces, analyzed the data and prepared the figures. J.M.S. also interpreted the measurements supported by J.O.W.. J.H. designed and validated the CRISPR/Cas9 constructs supported by J.M.S.. J.H. developed the cloning strategy, performed the molecular cloning and prepared the neuronal culture. J.O.W. reviewed the evaluation software for the MINFLUX traces. The widefield microscope setup was built by J.M.S. and J.E., who also wrote the software to control the setup. J.E. and E.D. provided technical supervision of the project. Developing MINFLUX for protein tracking, S.W.H. was responsible for overall project definition, supervision and steering. J.M.S., J.H. and S.W.H. wrote the manuscript with contributions from all authors.

## Acknowledgments

We thank Jana Kress, Birgit Koch, Paula Breuer and Alena Fischer for preparing the neuronal culture; Jessica Matthias for her initial support of the project and her introduction to the Abberior Instruments (AI) MINFLUX microscope and laboratory practice; Antonia Schach for her support with preliminary experiments and molecular cloning; Maria Augusta do Rego Barros Fernandes Lima and Lara Mayer for their initial assistance with the data analysis; Victor Macarrón Palacios for helping to maintain the MINFLUX microscope and advice on sample preparation; Tobias Engelhardt, Clara-Marie Gürth and Roman Schmidt (all of AI) for technical support; and our colleague Kai Johnsson for providing MaP555-Halo.

