## Supplements for "MINFLUX Reveals Dynein Stepping in Live Neurons"

### Supplementary Contents


1 Materials and Methods ... 3
1.1 CRISPR/Cas9 Knock-in Construct Design and Molecular Cloning ... 3
1.1.1 pORANGE-EGFP-Dync1i2 ... 3
1.1.2 pORANGE-HaloTag7-Dync1i2 ... 3
1.1.3 pORANGE-mEGFP-Dync1h1 ... 3
1.1.4 pORANGE-HaloTag7-Dync1h1 ... 4
1.1.5 CRISPR/Cas9 Knock-in Specificity Assessment ... 4
1.2 Sample Preparation ... 4
1.2.1 Primary Culture of Hippocampal Neurons and Electroporation ... 4
1.2.2 Fluorescent Labeling in Live Primary Neurons for MINFLUX Tracking ... 4
1.2.3 Fluorescent Labeling in Live Primary Neurons for Widefield Tracking ... 4
1.3 Microscope Setups ... 5
1.3.1 MINFLUX Setup ... 5
1.3.2 Widefield Setup ... 5
1.4 Dynein Tracking in Live Primary Neurons ... 5
1.4.1 MINFLUX Tracking Procedure ... 5
1.4.2 Widefield Measurements ... 6
1.5 Data Processing and Analysis ... 6
1.5.1 Analysis of MINFLUX Traces ... 6
1.5.2 Generation of Kymographs from Widefield Data ... 7
2 Supplementary Figures ... 8
3 Supplementary Tables ... 14
4 Supplementary Videos ... 19
5 Supplementary References ... 20

### 1 Materials and Methods

#### 1.1 CRISPR/Cas9 Knock-in Construct Design and Molecular Cloning

The general cloning strategy of the four plasmids used in this study, pORANGE-EGFP-Dync1i2, pORANGE-Halo7-Dync1i2, pORANGE-mEGFP-Dync1h1 and pORANGE-Halo7-Dync1h1 (Tab. S2), is described below. All plasmids were designed based on the CRISPR/Cas9 knock-in template vector pORANGE (gift from Harold MacGillavry, Addgene #131471) (1). GuideRNAs (gRNAs) near the 5′ end of dynein cytoplasmic 1 intermediate chain 2 (*Dync1i2*) exon 1 and dynein cytoplasmic 1 heavy chain 1 (*Dync1h1*) exon 1 were generated using the CRISPR design tool from Benchling based on scoring algorithms (2, 3) (Tab. S3, primers #1, #2, #9, #10). Target sequences and target sites are listed in Table S1.

##### 1.1.1 pORANGE-EGFP-Dync1i2

For plasmid construction, the pORANGE template vector was digested with BbsI (Thermo Fisher Scientific). Oligonucleotides with compatible BbsI overhangs (primers #1, #2, Tab. S3) were hybridized and ligated into the gRNA cassette of the digested pORANGE cloning template. The donor sequence containing EGFP flanked by two inverse gRNA sequences was generated stepwise. EGFP, linker sequences, and the 3′ inverse gRNA (downstream of EGFP) were generated by two consecutive PCRs (primers #3-6, Tab. S3). For the first PCR, primers #3 and #4 were used and pORANGE GFP-Actb KI #2 (gift from Harold MacGillavry, Addgene #139666) served as template to amplify EGFP and add linker sequences and restriction sites. For the second PCR, a different set of primers (primers #5, #6, Tab. S3) was used to amplify the PCR product of the first PCR, introducing more restriction sites. To insert EGFP, the second PCR product and the first pORANGE intermediate construct were digested with XbaI and BamHI (both Thermo Fisher Scientific) and ligated with T4 ligase. Finally, the 5′ inverse gRNAs (upstream of EGFP) with HindIII/NheI compatible ends (primers #7, #8, Tab. S3) were annealed and ligated into the HindIII and NheI (both Thermo Fisher Scientific) digested second intermediate construct. After bacterial transformation with the ligated plasmid, colonies containing the correct construct were identified by sequencing the purified plasmid (GeneJET Endo-free Plasmid Maxiprep Kit, Thermo Fisher Scientific, cat. K0861), which was later used for electroporation of primary hippocampal neurons.

##### 1.1.2 pORANGE-HaloTag7-Dync1i2

pORANGE-HaloTag7-Dync1i2 was generated by a subcloning step. The previous plasmid pORANGE-EGFP-Dync1i2 and an ordered HaloTag7 with corresponding restriction sites (Integrated DNA Technologies) were digested with NheI and KpnI (both NEB) to exchange the EGFP with the HaloTag7 fragment.

##### 1.1.3 pORANGE-mEGFP-Dync1h1

For the construction of the pORANGE-mEGFP-Dync1h1 vector, the first step was the same as for the plasmid construction of pORANGE-EGFP-Dync1i2. Different oligonucleotides (primers #9, #10, Tab. S3) were hybridized to target *Dync1h1* with the integration site in codon 1. The donor sequence containing mEGFP flanked by two inverse gRNA sequences was generated by PCR (primers #11, #12) and inserted into the pORANGE construct by restriction digestion with HindIII and MunI (both NEB), dephosphorylation and ligation. After bacterial transformation with the ligated plasmid, colonies containing the correct construct were identified by sequencing the purified plasmid.

##### 1.1.4 pORANGE-HaloTag7-Dync1h1

A subcloning step was used to generate pORANGE-HaloTag7-Dync1h1. The previous plasmid pORANGE-EGFP-Dync1h1 and an ordered HaloTag7 with corresponding restriction sites (Integrated DNA Technologies) were digested with NheI and SalI (both NEB) to exchange the mEGFP with the HaloTag7 fragment. The annotated map of the pORANGE-HaloTag7-Dync1h1 knock-in plasmid is shown in Fig. S1.

##### 1.1.5 CRISPR/Cas9 Knock-in Specificity Assessment

To assess the correct integration of respective tagging sequences into the genome cultured primary hippocampal neurons were electroporated with the following CRISPR/Cas9 knock-in plasmids: pO-EGFP-Dync1i2, pO-HaloTag7-Dync1i2, pO-EGFP-Dync1h1, pO-HaloTag7-Dync1h1. Genomic DNA was isolated using the GenElute Mammalian Genomic DNA Miniprep Kit (Sigma-Aldrich, cat. G1N70-1KT), followed by a PCR with respective primers. PCR products were purified after gel electrophoresis using the NucleoSpin Gel and PCR Clean Up kit (Macherey Nagel, cat. 740609) and sequenced (Eurofins Genomics). Correct incorporation of respective tagging sequences into the genome was confirmed using Geneious Prime (Biomatters, version 2023.2.1).

#### 1.2 Sample Preparation

##### 1.2.1 Primary Culture of Hippocampal Neurons and Electroporation

Primary hippocampal cultures were obtained from Wistar rats at postnatal day P0 to P1, as described previously (4). Prior to dissection, Ø 18 mm autoclaved coverslips were coated with 50 µL gold nanoparticle solution (BBI Solutions, cat. GC150), which was centrifuged down and diluted to 150 µL in PBS per coverslip. After at least 3 h, the remaining liquid was aspirated. The coverslips were then coated with 100 µg/mL poly-L-ornithine (Sigma-Aldrich, cat. P3655) and 1 µg/mL laminin (Corning, cat. 354232). 200,000 primary hippocampal neurons per coverslip were electroporated with 200 ng CRISPR/Cas9 knock-in plasmid using the Neon transfection system following manufacturer's instructions (Thermo Fisher Scientific, cat. MPK5000 & MPK10096) at 1,400 V and 3 pulses of 10 ms duration. Primary hippocampal neurons were maintained in an incubator (37 °C, 5% $CO_2$, 95% rH) in Neurobasal medium (Thermo Fisher Scientific, cat. 21103049) supplemented with 1% GlutaMAX (Thermo Fisher Scientific, cat. 35050061), 1% penicillin/streptomycin (Thermo Fisher Scientific, cat. 15070063) and 2% B27 (Thermo Fisher Scientific, cat. 17504044). On the first day in culture, the medium was replaced with fresh, supplemented Neurobasal medium.

##### 1.2.2 Fluorescent Labeling in Live Primary Neurons for MINFLUX Tracking

Just before the tracking experiment, primary hippocampal neurons were incubated with 100 pM JFX650 HaloTag Ligand (Promega, cat. CS315109) for 15 min in preconditioned medium and washed once with supplemented Neurobasal. Then, primary hippocampal neurons were simultaneously incubated with 100 nM MaP555-Halo (5) (kind gift from Kai Johnsson, MPIMR) and 1:20,000 BioTracker 488 Green Microtubule Cytoskeleton Dye (Sigma-Aldrich, cat. SCT142) for 15 min in preconditioned medium, washed once, followed by a 30 min washout in preconditioned medium. Coverslips were washed twice in ACSF buffer (27 mM Hepes pH 7.4, 126 mM NaCl, 2.5 mM KCl, 2.5 mM $CaCl_2$, 1.3 mM $MgCl_2$, 30 mM glucose) and transferred into a magnetic imaging chamber (Live Cell Instrument, Chamlide CMB) filled with ACSF for imaging.

##### 1.2.3 Fluorescent Labeling in Live Primary Neurons for Widefield Tracking

A similar sample preparation was performed for the widefield tracking experiments using coverslips without gold nanoparticles. Primary hippocampal neurons were simultaneously incubated with 10 nM MaP555-Halo (5) (kind gift from Prof. Kai Johnsson, MPIMR) and 1:40,000 BioTracker 488 Green Microtubule Cytoskeleton Dye (Sigma-Aldrich, cat. SCT142) for 15 min in preconditioned medium.

#### 1.3 Microscope Setups

##### 1.3.1 MINFLUX Setup

MINFLUX tracking experiments were performed with a commercially available MINFLUX system (Abberior Instruments, Göttingen, Germany) (6) built around a motorized inverted microscope IX83 (Olympus) with an 100x/1.4 objective lens (Olympus, UPLSAPO100XO). The microscope is equipped with 640 nm, 561 nm and 488 nm excitation laser lines and an LED lamp (Excelitas, XT720L) with corresponding filter sets. MINFLUX detection is performed with two avalanche photodiodes in the spectral ranges of 650-685 nm and 685-720 nm (photon counts were summed up). Confocal images are detected in the spectral ranges of 500-550 nm and 580-630 nm. Active stabilization of the sample is achieved by the reflection image of the gold nanoparticles fixed on the coverslip, resulting in a compensated drift of typically less than 1 nm (mean square displacement) in all directions. The setup is controlled via the Imspector software (Abberior Instruments, version 16.3.15631-m2205).

##### 1.3.2 Widefield Setup

A custom-built widefield microscope was set up for the widefield tracking experiments. The microscope is equipped with an inverted IX83 microscope body (Olympus), an 100x/1.4 objective lens (Olympus, UPLSAPO100XO) and a z-drift compensator unit (Olympus, IX3-ZDC). The setup has four excitation lasers with wavelengths of 640 nm (HÜBNER Photonics, 05-01 Cobolt Bolero), 570 nm (MPB Communications), 532 nm (Laser Quantum, gem 532) and 488 nm (Oxxius, LBX-488), whose intensity is controlled via an acousto-optic tunable filter (AA Opto Electronic, nC-TN-1001). For the activation of fluorophores, two lasers with wavelengths of 375 nm (Coherent, CUBE 375-16C) and 405 nm (Oxxius, LBX-405) can be used. Two quad-band filter sets are available for multiplex imaging, one for alternating excitation of 405 nm, 488 nm, 532 nm and 640 nm (AHF analysentechnik, F72-866 and F73-866S) and the other for alternating excitation of 405 nm, 488 nm, 570 nm and 640 nm (AHF analysentechnik, F72-832 and F73-832S). When the eyepieces are used, the sample is illuminated with an LED lamp (Excelitas, XT720L) with corresponding filter sets. Emitted light is directed through a 1.6x magnification onto an EMCCD camera (Oxford Instruments, Andor iXon$^{EM}$+ DU-897D-CSO-BV), resulting in an effective pixel size of 100 nm and a field of view of 51.2 x 51.2 µm$^2$. The beam angle is controlled via a motorized stage enabling total internal reflection fluorescence (TIRF) illumination. A custom-written LabVIEW program (National Instruments, version 20.0.1f1) controls all components of the setup.

#### 1.4 Dynein Tracking in Live Primary Neurons

After six to nine days in culture, a fresh sample was prepared for each of the tracking measurements. Using dynein labeled with Map555-Halo or tagged with EGFP, a CRISPR/Cas9 knock-in positive neuron was identified with the eyepieces and LED illumination. Axons were identified by morphology as the longest and smoothest processes. In cases where the axon was not clearly identifiable, a different cell was sought.

##### 1.4.1 MINFLUX Tracking Procedure

Several regions of interest (ROI) along the axon were successively selected for MINFLUX tracking. To achieve a temporal resolution of less than 1 ms, the pattern dwell time was set to 600 µs and the laser power of the 640 nm laser line was set to 135 µW (measured at the periscope) to achieve photon rates above 125 kHz. The pinhole was set to 0.93 AU. During the MINFLUX measurement, the ROI is scanned and as soon as the defined photon threshold is exceeded, the specified iterative MINFLUX localization procedure (Tab. S4) is started. The diameter L of the targeted coordinate pattern (TCP) is gradually reduced and the laser power is successively increased. The MINFLUX localization pattern remains in the last iteration step and tracks the emitter position with the donut-shaped excitation beam until the detection signal is lost or the maximum center frequency ratio (CFR, Tab. S4) (6) is exceeded, resulting in a trace of successive localizations.

##### 1.4.2 Widefield Measurements

For widefield tracking, ROIs (51.2 x 51.2 $\mu m^2$) were successively selected along the axon. In each ROI, stained microtubules were imaged and bleached with the 488 nm laser at a focal intensity of 92 $W/cm^2$ and near TIRF conditions. Subsequently, videos of at least 1 min were recorded in the same ROI with the 532 nm laser at a focal intensity of 92 $W/cm^2$ and an exposure time of 100 ms.

#### 1.5 Data Processing and Analysis

In all presented figures, n denotes the number of data points and N denotes the number of independent experiments.

##### 1.5.1 Analysis of MINFLUX Traces

Custom written MATLAB scripts (MathWorks, version R2022a) were used to analyze the MINFLUX traces and prepare the resulting figures. Four filter criteria were used to pre-filter processive traces from the dataset. First, traces with fewer than 200 localizations were filtered out. In the second filtering step, traces with a photon rate above 300 kHz were excluded. From the remaining traces, the main propagation direction (on-axis) and its perpendicular direction (off-axis) were determined using singular value decomposition with the MATLAB function *svd* (7). The traces were then transformed into the new on-/off-axis coordinate system. In the third filtering step, traces whose singular value $\sigma_1$ was less than 20 nm were discarded, as these traces do not show processive motion. In the fourth filter step, traces with similar variance in both axes ($\sigma_1/\sigma_2 < 3$) were discarded. The steps of the traces were identified with the AutoStepfinder algorithm (8) and then divided into retrograde and anterograde segments (at least 50 nm processive runs). The trace segments between consecutive runs in opposite directions was defined as a direction reversal. Pauses within processive runs were defined as trace segments where the speed was less than 100 nm $s^{-1}$ for at least 100 ms. Three models were fitted to the dwell time distributions using maximum likelihood estimates with the MATLAB function *mle*, a single exponential ($ke^{-k\tau}$), a convolution of two exponentials with the same rate ($k^2\tau e^{-k\tau}$), and a convolution of two exponentials with different rates ($k_1k_2/(k_1 - k_2)(e^{-k_2\tau} - e^{-k_1\tau})$). The model that best describes the data was determined using Akaike weights as a measure of the probability that one of the candidate models is the best model given the data and the three candidate models (9). Note that the Akaike Information Criterion ($\mathrm{AIC} = 2k - 2\ln(\hat{L})$), where $k$ is the number of estimated parameters in the model and $\hat{L}$ is the likelihood function, penalizes the model with two different rate constants more than the other two models. The localization precision of a trace was calculated from the difference of successive localizations as described previously (10). The step detection uncertainty was estimated based on step size and plateau length (8).

##### 1.5.2 Generation of Kymographs from Widefield Data

Kymographs were generated from the recorded widefield videos along the axon using the Fiji ImageJ (11) macro KymographClear (12). Segments of anterograde and retrograde movement were manually traced and exported using the KymographDirect program (12). The exported data were visualized using MATLAB. Only segments with a run length longer than 1 µm were considered.

#### 2 Supplementary Figures

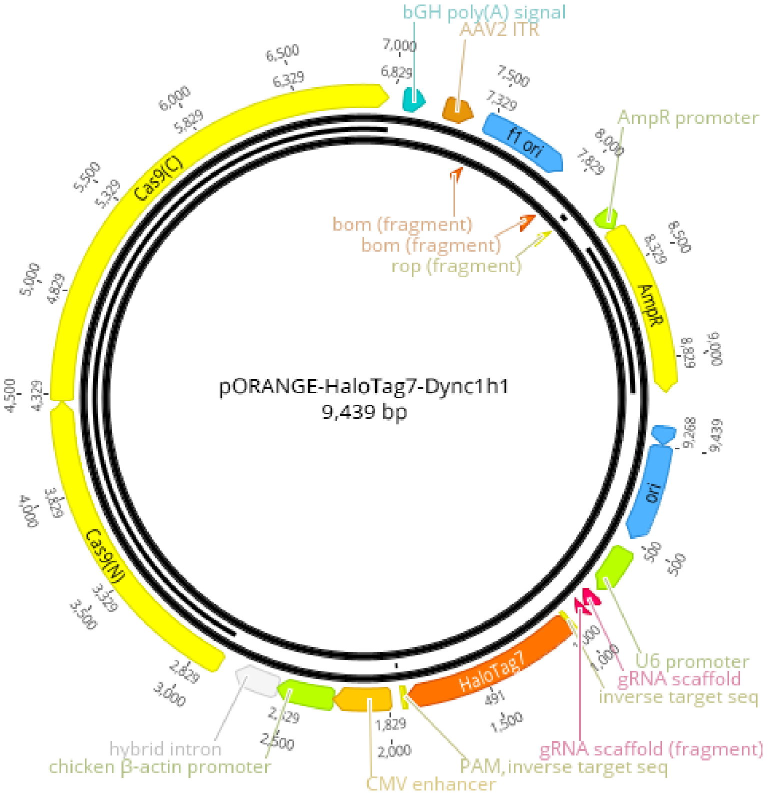


**Figure S1. Plasmid map of pORANGE-HaloTag7-Dync1h1.** The plasmid shown here is used to knock-in a HaloTag7 sequence at the 5′ end of the rat gene *Dync1h1*, resulting in the N-terminus of endogenous DHC being tagged with a HaloTag7. The three other plasmids generated are similar to the one shown here.

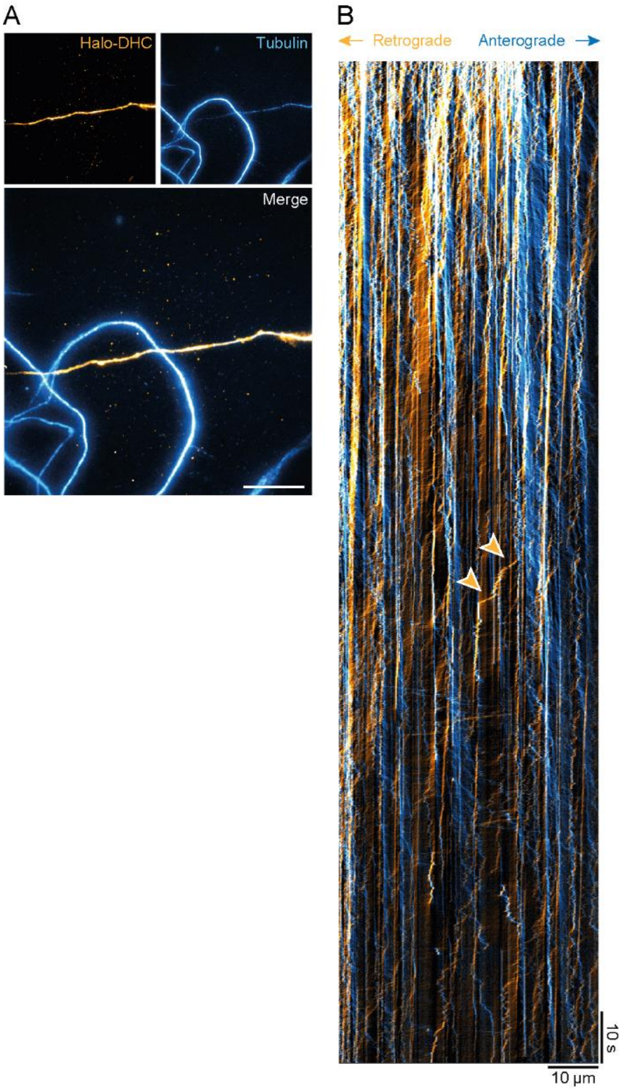


**Figure S2. Example widefield acquisition and kymograph of Halo-DHC.** (A) Widefield acquisition of an axon with Halo-DHC particles labeled with 10 nM MaP555-Halo and stained microtubules. (B) The kymograph showing a retrogradely moving Halo-DHC particle with two fast runs marked with arrows. This corresponds to video S1. Scale bar: 10 µm (A).

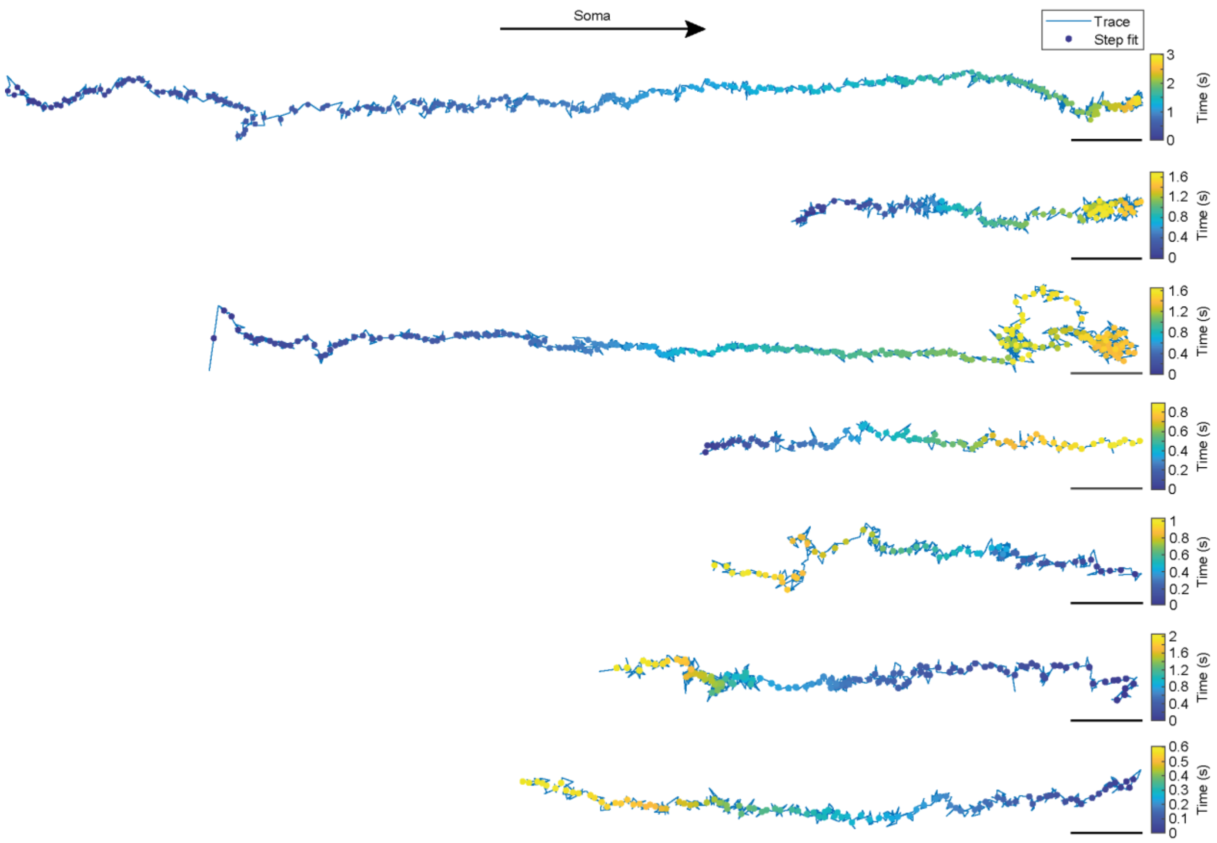


**Figure S3. Example MINFLUX traces of endogenous dynein in live primary neurons.** These raw traces (Halo-DHC and Halo-DIC) shown here also contain steps that were filtered out as pauses or reversals in the subsequent segmentation analysis, since the step finding algorithm used fits steps to the entire traces including pauses or reversals. Scale bars: 100 nm.

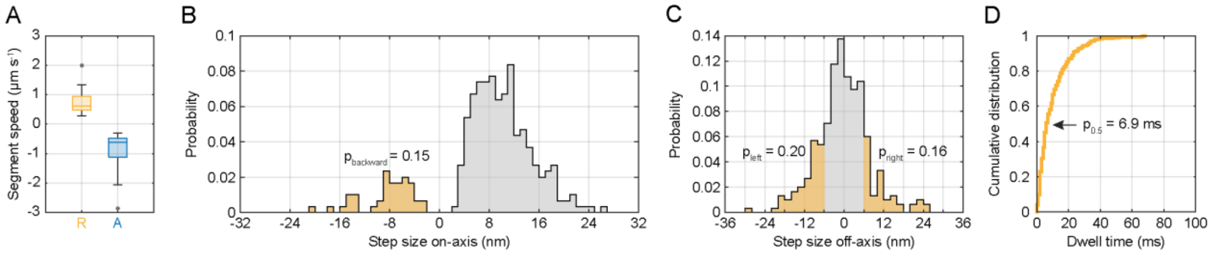


**Figure S4. Stepping behavior of dynein intermediate chain in live neurons.** (A) Boxplot of segment speeds in the retrograde (n = 20) and anterograde directions (n = 26) of Halo-DIC. (B) On-axis step size distribution of Halo-DIC with indicated proportion of backward steps (n = 298, N = 12). (C) Off-axis step size distribution of Halo-DIC with highlighted proportions of off-axis steps larger than 6 nm in either direction (n = 298, N = 12). (D) Cumulative distribution of dwell times between consecutive steps of Halo-DIC (n = 278, N = 12).

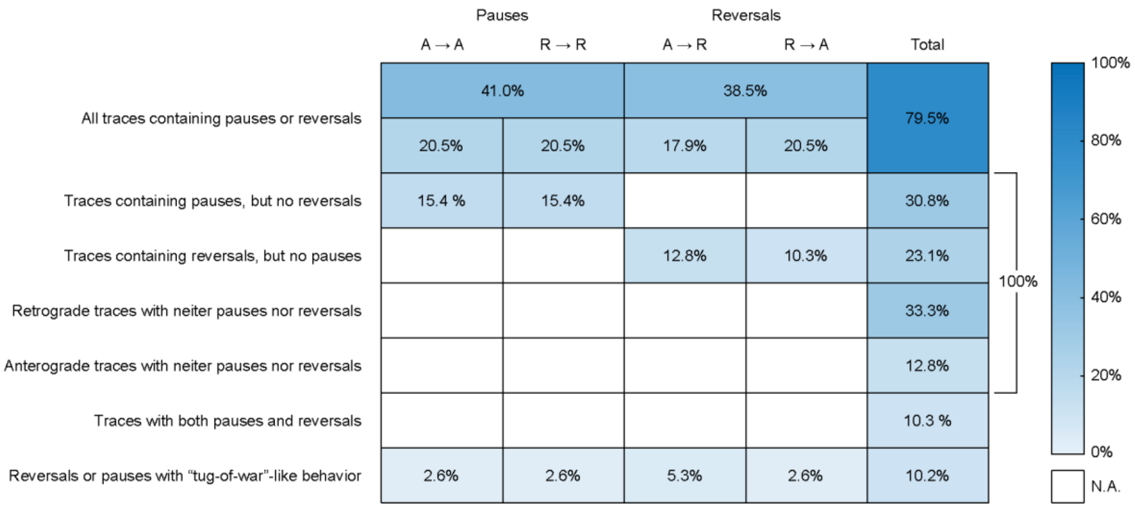


**Figure S5. Overview of different scenarios of pauses and reversals.** MINFLUX traces were classified into processive anterograde and retrograde trace segments (on-axis displacement was at least 50 nm). Segments between consecutive runs in opposite directions were defined as a direction reversal (n = 24). Pauses within consecutive runs were defined as trace segments where the speed was less than 100 nm $s^{-1}$ for at least 100 ms (n = 25). It should be noted that traces in the first two rows may be counted twice if they exhibited both pauses and reversals.

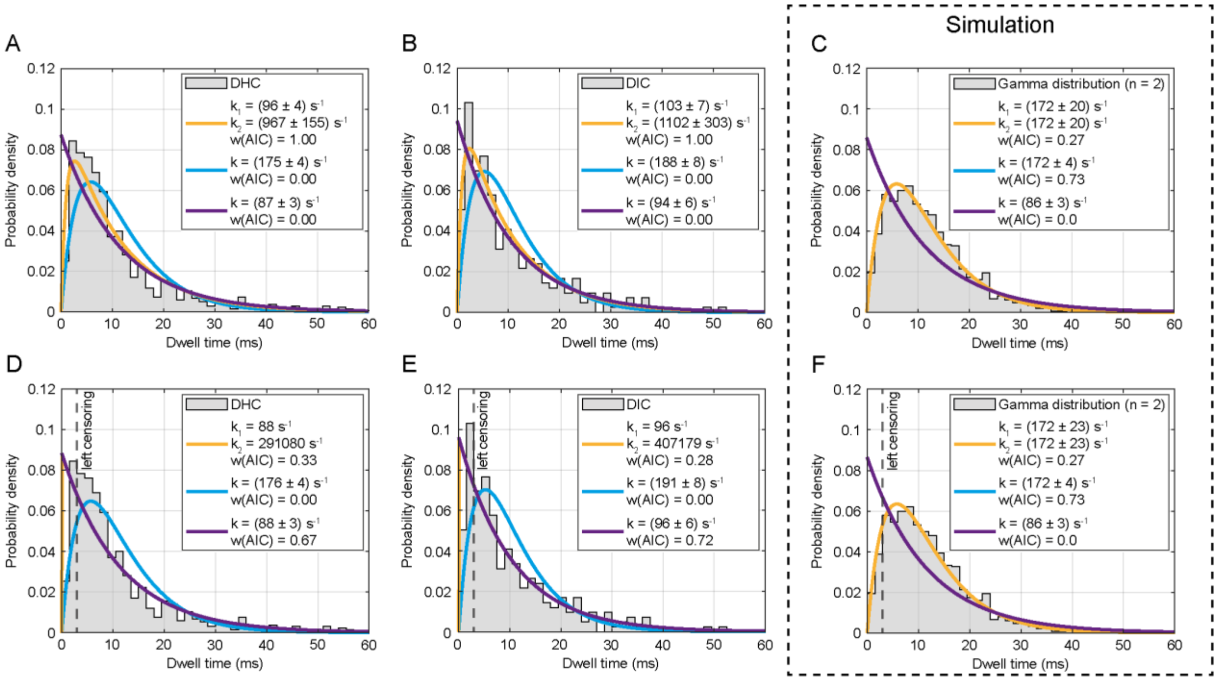


**Figure S6. Dwell time distributions of endogenous dynein with left censoring compared to simulated gamma distribution.** (A) Uncensored dwell time distribution of DHC (n = 900, N = 6) is best described by a convolution of two exponentials with two different rates $(k_1k_2/(k_1 - k_2)(e^{-k_2\tau} - e^{-k_1\tau}))$ (orange curve), suggesting that dynein stepping is based on one rate-limiting process $k_1$ and a faster process $k_2$. (B) Uncensored dwell time distribution of DIC (n = 278, N = 17) leads to the same considerations as for DHC. (C) Simulated gamma distribution of degree n = 2 represents a process that is described by two consecutive events (n = 1000), i.e. the case when dynein stepping would be based on the consecutive consumption of two ATP per step. In this case, the distribution is best described by the cyan curve, which fits a convolution of two exponentials with the same rate constant ($k^2\tau e^{-k\tau}$) to the distribution. The orange curve overlaps exactly with the cyan curve, whose fit parameters are in this case the same as those of the cyan curve ($k_1 = k_2 = k$). (D) Left censoring of the DHC dwell time distribution corresponds to the case where it is assumed that extremely fast steps ($\tau < 3$ ms) are largely missed and therefore underrepresented. Consequently, data points with values less than 3 ms are not considered by the fitting algorithm. In this case, the distribution is best described by the purple curve, which corresponds to a single exponential ($ke^{-k\tau}$), i.e. dynein stepping is based on a single rate-limiting process (one ATP per step). The orange curve, representing a convolution of two exponentials with unequal rate constants, overlaps exactly with the purple curve, and the low rate constant $k_1$ corresponds to that of the single exponential model, while the high rate constant $k_2$ becomes infinite. 148 data points were excluded by the left censoring. (E) Left censoring of the DIC dwell time distribution leads to the same considerations as for DHC. 64 data points were excluded by the left censoring. (F) Left censoring of the simulated gamma distribution does not influence the fit in contrast to the measured dwell times. 87 data points were excluded by left censoring. All panels show fit parameters and corresponding Akaike weights of the three assumed models, a convolution of two exponential decays with two equal rate constants (cyan) and two unequal rate constants (orange), or a single exponential decay (purple).

### 3 Supplementary Tables

**Table S1**. **Target sequences and target sites for knock-in at rat *Dync1i2* exon 1 and *Dync1h1* exon 1.**

| **Gene** | **Name** | **Target sequence (PAM is underlined)** | **Site of integration (in amino acid)** |
|---|---|---|---|
| *Dync1i2* | dynein cytoplasmic 1 intermediate chain 2 | TAATTCACTTTTGTCTGACATGG | S2 |
| *Dync1h1* | dynein cytoplasmic 1 heavy chain 1 | CTTCCGCGGATCCGCCATGTCGG | M1 |

**Table S2. Overview of generated CRISPR/Cas9 knock-in plasmids for endogenous tagging of dynein.**

| Plasmid | Purpose |
|---|---|
| pO-EGFP-Dync1i2 | Endogenous N-terminal tagging of dynein intermediate chain with EGFP (ORANGE-based knock-in) |
| pO-HaloTag7-Dync1i2 | Endogenous N-terminal tagging of dynein intermediate chain with HaloTag7 (ORANGE-based knock-in) |
| pO-EGFP-Dync1h1 | Endogenous N-terminal tagging of dynein heavy chain with EGFP (ORANGE-based knock-in) |
| pO-HaloTag7-Dync1h1 | Endogenous N-terminal tagging of dynein heavy chain with HaloTag7 (ORANGE-based knock-in) |

**Table S3**. **Primers for CRISPR/Cas9 knock-in plasmid generation.**

| No. | Primer name | Sequence 5´ - 3´ | Purpose |
|---|---|---|---|
| **#1** | ORANGE_Dync1i2-KI-gRNA1-fw | CACCGTAATTCACTTTTGTCTGACA | gRNA for tagging Dync1i2 |
| **#2** | ORANGE_Dync1i2-KI-gRNA1-rev | AAACTGTCAGACAAAAGTGAATTAC | gRNA for tagging Dync1i2 |
| **#3** | PCR_fw_EGFP_HindIII_NheI | CTCTAGAAGCTTTCGAACCATGGCTAGCGGAGTGAGCAAGGGCGAGGAGCTGTTCACC | Forward primer for 1. PCR to amplify insert EGFP |
| **#4** | PCR_rev_XhoI gRNADyn EGFP | GTACATCTCGAGGAGAAGACCCCGAGCCATGTCAGACAAAAGTGAATTACCAGCGCTTCCACTCTTGTACAGCTCGTCCATGCCGAGAGTGAT | Reverse primer for 1. PCR to amplify insert EGFP (Addgene #139666) |
| **#5** | PCR_fw_XbaI_HindII_NheI | GTCACGGTCTAGAAGCTTTCGAACCATGGCTAGC | Forward Primer for 2. PCR to amplify PCR product 1 |
| **#6** | PCR_rev_BamHI_XhoI | GTACATGGATCCATCTCGAGGAGAAGACCCCGAGCCATG | Reverse Primer for 2. PCR to amplify PCR product 1 |
| **#7** | gRNA_Dync1i2_HindIII_NheI_fw | AGCTTTCGAAGACCCTAGATAATTCACTTTTGTCTGACATGGGCTTCGCCATGG | Inverse gRNA to cut out EGFP |
| **#8** | gRNA_Dync1i2_HindIII_NheI_rev | CTAGCCATGGCGAAGCCCATGTCAGACAAAAGTGAATTATCTAGGGTCTTCGAA | Inverse gRNA to cut out EGFP |
| **#9** | ORANGE_Dync1h-KI-gRNA1-fw | CACCGCTTCCGCGGATCCGCCATGT | gRNA for tagging of Dync1h1 |
| **#10** | ORANGE_Dync1h1-KI-gRNA1-rev | AAACACATGGCGGATCCGCGGAAGC | gRNA for tagging of Dync1h1 |
| **#11** | Restriction_fw mEGFP_Dyn1h1 | CTGCAGACAAATGGCTCTAGAAGCTTCCGACATGGCGGATCCGCGGAAGGCTAGCATGGTGAGCAAGGGCGAG | PCR forward primer to amplify insert mEGFP |
| **#12** | Restriction_rev mEGFP_Dyn1h1 | GACGCGTCCTAGGATCCTCGAGTCGACAATTGCTTCCGCGGATCCGCCATGTCGGAACCCTTGTACAGCTCGTCCATGCC | PCR reverse primer to amplify insert mEGFP |

**Table S4. MINFLUX tracking sequence.** The minimum number of detected photons (photon count) refers to the total dwell time of the entire pattern (e.g. 600 µs for final iteration). If the minimum number of photons is not reached during a pattern, the entire pattern is repeated to reach the minimum number of photons. This increases the time interval to the next localization. L, pattern diameter; CFR, center frequency ratio; TCP, targeted coordinate pattern.

| **Iteration** | **1** | **2** | **3** | **4** |
|---|---|---|---|---|
| **L (nm)** | 284 | 302 | 151 | 75 |
| **Single pattern dwell time (µs)** | 50 | 50 | 50 | 200 |
| **Pattern repeat** | 1 | 1 | 1 | 3 |
| **Minimum photon counts** | 40 | 20 | 10 | 75 |
| **Background threshold (kHz)** | 15 | 30 | 30 | 40 |
| **Maximum CFR** | n.a. | 0.5 | n.a. | 0.8 |
| **Laser power multiplication** | 1 | 1 | 2 | 3 |
| **TCP mode** | Prescan | Hexagon | Hexagon | Hexagon |

**Table S5. Overview of various parameters of endogenous dynein stepping.** The step size parameters were calculated from all retrograde segments of recorded MINFLUX traces. Pause and direction reversal durations were calculated from all MINFLUX traces (Halo-DHC and Halo-DIC combined). MAD, median absolute deviation; SD, standard deviation.

| **Parameter** | **Quantity (median ± MAD)** | | **Quantity (mean ± SD)** | |
|---|---|---|---|---|
| | **Halo-DHC** | **Halo-DIC** | **Halo-DHC** | **Halo-DIC** |
| Overall step size | (7.9 ± 2.3) nm | (8.6 ± 3.5) nm | (8.0 ± 5.0) nm | (7.6 ± 8.0) nm |
| Forward step size | (8.0 ± 2.2) nm | (9.8 ± 3.1) nm | (8.7 ± 3.8) nm | (10.4 ± 4.6) nm |
| Backward step size | (5.7 ± 2.0) nm | (7.4 ± 2.1) nm | (7.2 ± 5.2) nm | (8.3 ± 4.1) nm |
| Segment speed | (670 ± 340) nm $s^{-1}$ | (620 ± 180) nm $s^{-1}$ | (660 ± 370) nm $s^{-1}$ | (740 ± 420) nm $s^{-1}$ |
| Pause duration (A → A) | (166 ± 50) ms | | (232 ± 180) ms | |
| Pause duration (R → R) | (141 ± 16) ms | | (183 ± 104) ms | |
| Reversal duration (A → R) | (11 ± 6) ms | | (41 ± 58) ms | |
| Reversal duration (R → A) | (20 ± 11) ms | | (67 ± 104) ms | |

### 4 Supplementary Videos

**Video S1.** The video shows a widefield acquisition of an axon with Halo-DHC particles labeled with 10 nM MaP555-Halo. The playback speed is 10-fold. A retrogradely moving Halo-DHC particle with two fast runs is marked with an orange arrowhead (visible in the middle of the video). The white arrow indicates the direction of the soma. Scale bar: 10 µm.
